# Filament mechanics and turnover organize dynamic load bearing structures in a cellular-scale model of the actomyosin cortex

**DOI:** 10.64898/2026.11.10.765540

**Authors:** Kohsuke Tsukui, Hiromi Miyoshi, Naoya Sakamoto, Hiro Wakimura, II Satoshi

## Abstract

The actomyosin cortex controls cell shape and movement by generating and transmitting contractile forces. Its actin filament (AF) network adopts diverse spatial organizations, while spatially biased contractility drives cortical flow that transports components and reorganizes the network. However, how turnover and spatial redistribution organize and maintain tension-bearing structures in a dynamic steady state, and how localized contractility reorganizes them through flow, remain unclear. We developed a cellular-scale discrete model of the actomyosin cortex representing AFs, cross-linkers, and myosin minifilaments. Local surface stress was reconstructed from discrete-element internal forces based on continuum mechanics. We related cortical tension to component abundance, tensile deformation, AF orientation, turnover, and network connectivity, and compared different AF disassembly, relocation, and myosin-localization mechanisms. Simulations produced a heterogeneous, network-like distribution of local cortical tension. Tensile energy per element correlated more strongly with local tension than component abundance, with the strongest correlation for AFs. Highly stretched AFs preferentially occurred in high-tension regions and aligned with the first-principal-tension orientation, indicating anisotropic load-bearing pathways. Despite continuous component replacement and reorganization of local tension patterns, global AF-network connectivity and cell-scale cortical tension were maintained. The characteristic time of tension-pattern reorganization was closest to the persistence time of myosin, suggesting an association between myosin dynamics and rapid force redistribution. AF relocation to low-density regions replenished spatial gaps, whereas bending-dependent disassembly limited the accumulation of strongly deformed AFs. Direct myosin localization and AF accumulation produced comparable equatorward cortical flows but differed in the spatial organization of the tension-bearing network. Collectively, these results demonstrate that cellular-scale cortical tension is organized not only by component abundance but also by filament-scale mechanical loading, AF orientation, and network connectivity.

## 1 Introduction

The actomyosin cortex is a thin cytoskeletal layer formed immediately beneath the plasma membrane and is composed of actin filaments (AFs), nonmuscle myosin II, and actin-binding proteins, including cross-linkers (CLs; e.g., *α*-actinin). Cortical tension is one of the principal mechanical properties of the actomyosin cortex, and its spatial and temporal regulation is essential for the maintenance and deformation of cell shape, cell migration, and cell division (Salbreux et al., 2012; Kelkar et al., 2020). Within the cortex, nonmuscle myosin II assembles into bipolar minifilaments whose motor heads bind to separate AFs and generate relative sliding between them (Murrell et al., 2015; Chinthalapudi and Heissler, 2024). Depending on the geometric arrangement, deformation, and mechanical connectivity of the AFs, this local myosin activity generates contractile stress within the network (Lenz, 2014; Murrell et al., 2015). The resulting stress is transmitted through the AF network across the cortical surface and manifests as cortical tension at the cellular scale. Understanding the mechanical basis of cortical tension therefore requires not only characterization of local force generation by myosin but also clarification of the AF network structure that bears and transmits these forces.

The cortical AF network adopts different spatial organizations depending on cell type and physiological state, including meshworks, branched networks, and filament bundles (Svitkina, 2020). Mathematical models based on local interactions among cortical components have reproduced AF network formation and periodic reorganization (Hernández-del Valle et al., 2022), as well as the spontaneous emergence of spatiotemporal patterns such as localized clusters, contractile pulses, and network-like structures (Staddon et al., 2022). The spatial distribution and directionality of cortical tension also vary with cellular state. In-plane anisotropy of cortical tension has been observed during cell polarization (Mayer et al., 2010), and the relationship between in-plane AF orientation and direction-dependent cortical tension has been examined during cytokinesis (Spira et al., 2017). Moreover, network compression caused by cortical flow has been shown to contribute to the alignment of AFs during cytokinesis (Reymann et al., 2016). These studies have characterized cellular-scale AF network reorganization and the associated spatial and directional variations in cortical tension. At more local scales on the cell surface, however, the relationships of AF distribution, mechanical loading, and orientation with the local stress and tension state remain insufficiently characterized.

The spatial organization of the cortex is not static but is continuously remodeled through AF polymerization, depoly-merization, and severing, as well as through the turnover of myosin and CLs (Salbreux et al., 2012; Kelkar et al., 2020). As the principal structural component of the network, AFs are renewed not only in quantity but also in spatial organization. In reconstituted entangled F-actin networks, cofilin-dependent severing promotes rapid AF turnover and network reorganization (McCall et al., 2019). Experiments on single AFs have shown that tension suppresses cofilin-dependent severing, whereas experiments on AF bundles have shown that tension reduces the frequency of cofilin binding (Hayakawa et al., 2011). A mesoscale model of cofilin-bound AFs predicted that compressive buckling promotes cofilin-dependent severing, whereas torsional deformation preferentially promotes cofilin dissociation rather than AF severing (Schramm et al., 2017). A subsequent study predicted that, compared with axial extension, bending and torsion produce more heterogeneous interfacial strain and partial bond rupture, thereby increasing the susceptibility of AFs to fragmentation (Schramm et al., 2019). Regarding AF replenishment, a two-dimensional agent-based model demonstrated that turnover rules pre-scribing the position and orientation of newly introduced AFs differentially affect bundle formation and sustained contractility (Tam et al., 2026). Collectively, these findings indicate that AF network remodeling depends not only on filament exchange itself but also on the mechanical state of the AFs being removed and the spatial rule by which new AFs are introduced. The structures that bear cortical tension should therefore be viewed not as fixed AF arrangements but as dynamic load-bearing structures that are continually renewed through AF removal and replenishment.

These observations raise the question of how tension-bearing actomyosin networks are mechanically organized by local interactions and turnover, and how their load-bearing function is maintained in a dynamic steady state despite continuous component exchange. Mathematical models have examined stress generation, dissipation, and long-range flow regulated by AF turnover and CL dynamics (McFadden et al., 2017); bundle formation, contraction, and polarity alignment arising from the coupling of component kinetics and mechanical deformation (Popov et al., 2016); and global contraction patterns governed by the interplay among network percolation, advection, turnover, and system size (Krishna et al., 2024). These studies provide complementary insights into how network mechanics and component dynamics influence network organization and contractile behavior. Building on these findings, further investigation is needed to clarify how the spatial distribution and reorganization of AF load-bearing states across the entire cortical surface relate to the maintenance of global network connectivity and cell-scale cortical tension.

In this study, we aimed to determine how the network structure bearing cortical tension is mechanically organized in a dynamic steady state and maintained under continuous component exchange at the cellular scale. To this end, we developed a discrete model of the actomyosin cortex that explicitly represents AFs, CLs, and myosin minifilaments as mechanical elements and relates the mechanical state of individual AFs to the local stress on the cell surface. Using this model, we analyzed the relationships of local AF abundance, mechanical loading, and orientation with the local stress distribution, as well as the roles of AF replenishment and mechanically regulated removal in maintaining load-bearing structures and cellular-scale cortical tension. Finally, we examined how two idealized modes of equatorial cortex organization, direct localization of myosin on the existing AF network and localization mediated by AF accumulation, differently affect the organization of load-bearing structures, cortical tension, and cell-scale cortical flow.

## 2. Methods

### 2.1 Discrete model of the actomyosin cortex

Figure 1 illustrates the present model. We assumed a thin cortical layer beneath a fixed spherical plasma membrane. The cortical region was defined as a spherical shell with cell radius *R*_*c*_ and cortical thickness *t*_*c*_. The cortex was represented as a discrete network consisting of AFs, CLs, and myosin minifilaments (Figs. 1(A) and (B)). Each AF was modeled as a discrete fiber capable of stretching and bending and was composed of multiple nodes connected by stretching line elements. Each CL and myosin minifilament was modeled as a stretching line element that mechanically connected a pair of AFs. The plasma membrane was treated as a fixed spherical boundary and was not allowed to deform or move. To represent ERM-mediated anchoring of the cortical AF network to the plasma membrane without explicitly modeling ERM proteins, selected AF nodes near the membrane were constrained to move tangentially along the fixed spherical surface.

**Fig. 1.**
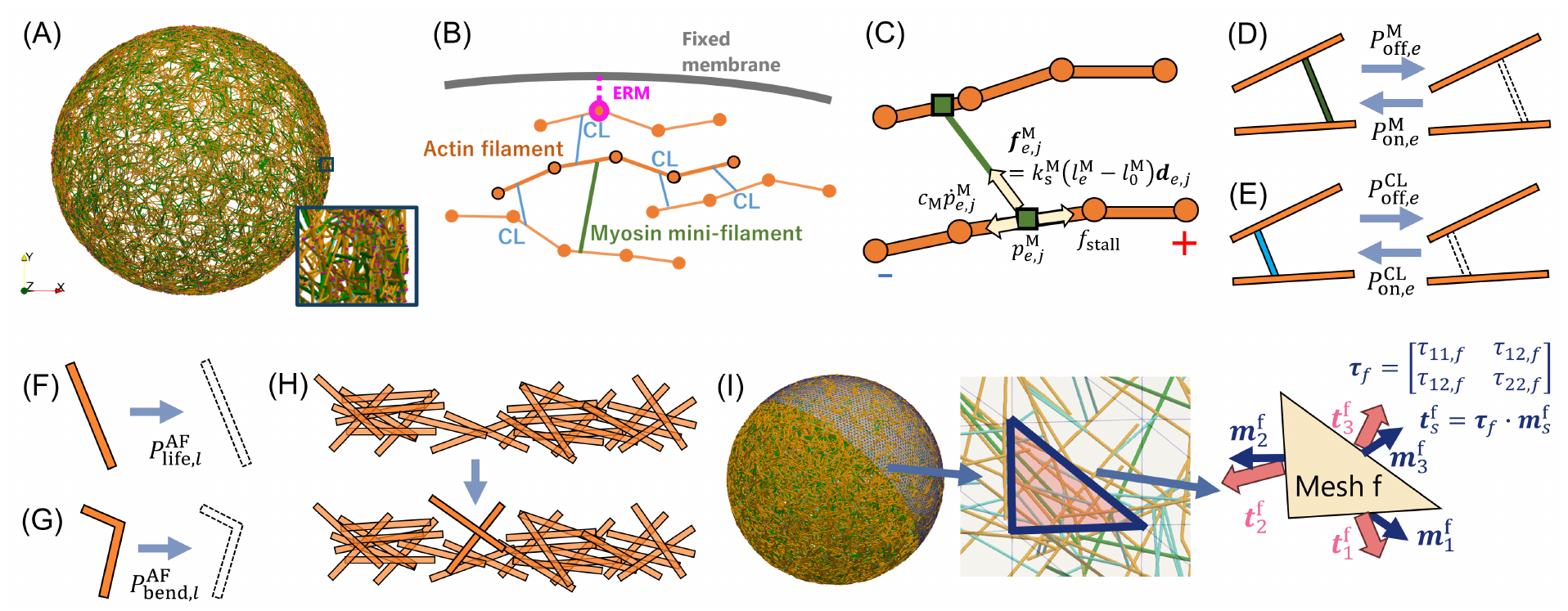
Overview of the discrete actomyosin cortex model. (A) Spherical-shell network of AFs (orange), CLs (light blue), and myosin minifilaments (green) beneath a fixed spherical plasma membrane. (B) AFs represented by nodes and line elements and interconnected by CLs and myosin minifilaments. Selected AF nodes were tangentially constrained to represent ERM-mediated membrane anchoring. (C) Myosin minifilaments connecting two AFs, with attachment points moving toward the respective plus ends. (D) Stochastic binding and unbinding of myosin minifilaments and (E) CLs. (F) Constant-rate lifetime-dependent and (G) bending-dependent AF disassembly. (H) Immediate AF repositioning to low-density surface regions after disassembly. (I) Local surface stress tensor reconstructed from internal forces transmitted across mesh edges by least-squares fitting.

#### 2.1.1 Mechanical model and equation of motion for AFs

The motion of the *i*-th AF node was described by the following overdamped equation:

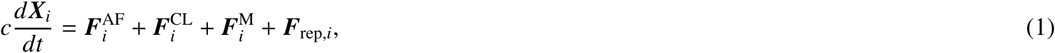

where ***X***_*i*_ is the position of the node, *c* is the drag coefficient associated with the surrounding medium, 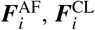, and 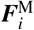 are the forces exerted by AFs, CLs, and myosin minifilaments, respectively, and ***F***_rep,*i*_ is the repulsive force exerted by the fixed spherical plasma-membrane boundary.

The stretching energy of line element *e* of component type *α* ∈ {AF, CL, M} was defined as

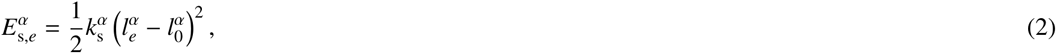

where 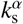 is the spring constant, and 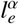 and 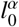 are the current and reference lengths of line element *e* of component type *α*, respectively.

The bending energy associated with internal node *q* of an AF, excluding the two terminal nodes, was defined as

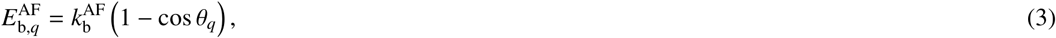

where 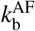is the bending stiffness and *θ*_*q*_ is the angle between the two AF line elements connected at node *q*.

Let *L*^*α*^ denote the set of line elements of component type *α* ∈ {AF, CL, M}, and let Q^AF^ denote the set of all internal AF nodes, excluding the two terminal nodes of each AF. The mechanical energy associated with component type *α* was defined as

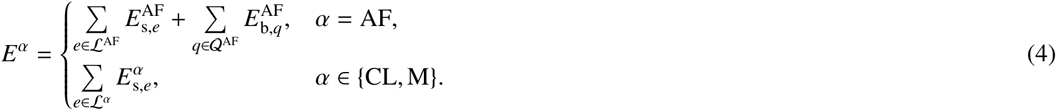

The force exerted by component type *α* on the *i*-th AF node was calculated as

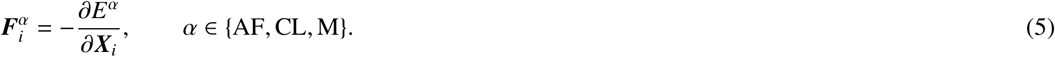

The short-range interaction between an AF node and the fixed plasma-membrane boundary was modeled using a truncated Morse interaction:

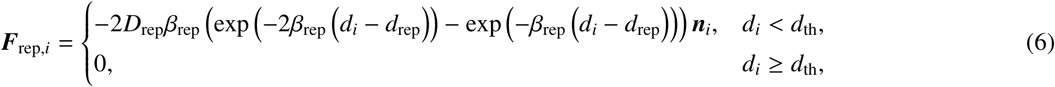

where *d*_*i*_ is the signed distance from the *i*-th AF node to the plasma-membrane surface and is defined as positive inside the spherical boundary. The unit vector ***n***_*i*_ points from the AF node toward the plasma-membrane surface. The parameters *D*_rep_, *β*_rep_, *d*_rep_, and *d*_th_ denote the interaction strength, inverse interaction length, equilibrium distance, and cutoff distance, respectively. In the present study, the cutoff distance was set equal to the equilibrium distance, *d*_th_ = *d*_rep_, so that only the repulsive branch of the Morse interaction was applied. The parameter values were *D*_rep_ = 10^−5^ J*/*m^2^, *β*_rep_ = 20 *µ*m^−1^, and *d*_rep_ = *d*_th_ = 0.05 nm.

For selected AF nodes representing ERM-mediated anchoring to the plasma membrane, the normal component of the nodal velocity was removed to constrain the nodes to tangential motion along the fixed spherical surface:

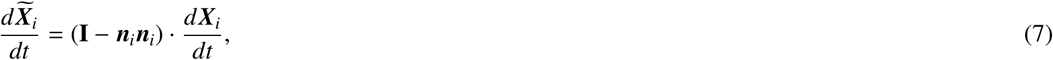

where **I** is the identity tensor and 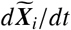 is the constrained tangential velocity.

#### 2.1.2. Dynamics and binding kinetics of myosin minifilaments and CLs

Each myosin minifilament was modeled as a stretching line element with two endpoints. While myosin element *e* was bound to two AFs, the motion of each endpoint *j* ∈ {1, 2} along its attached AF was governed by

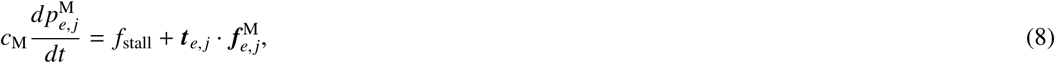

where 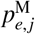is the attachment position of endpoint *j* of myosin element *e* on the AF, *c*_M_ is the friction coefficient for motion along the AF, *f*_stall_ is the stall force, and ***t***_*e, j*_ is the unit tangent vector directed toward the plus end of the AF line element to which the endpoint is attached. Here, 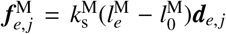 is the elastic force exerted by myosin element *e* at endpoint *j*, where ***d***_*e, j*_ is the unit vector directed from endpoint *j* toward the other endpoint of the myosin element. The polarity of each AF was prescribed, and each myosin endpoint moved toward the corresponding plus end. When either endpoint reached the plus end of its attached AF, the entire myosin element detached and subsequently underwent stochastic rebinding.

The binding and unbinding of myosin minifilaments and CLs were modeled as stochastic processes (Figs. 1(D) and (E)). For component type *α* ∈ {M, CL}, the binding and unbinding probabilities during a time step Δ*t* were given by

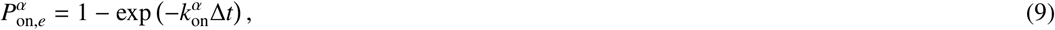

and

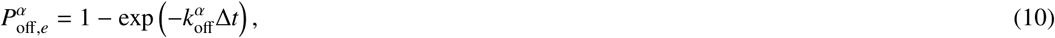

respectively, where 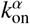 and 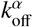 are the corresponding binding and unbinding rate constants. Myosin and CL elements were treated as either fully bound, with their two endpoints attached to two distinct AFs, or fully unbound; states with only one bound endpoint were not considered.

The same geometric procedure was used for the initial binding and subsequent rebinding of myosin and CL elements. First, an AF line element and a point on it were selected randomly as the first attachment site. Candidate second attachment sites were then identified at the intersections of line elements belonging to neighboring, distinct AFs with a sphere centered at the first attachment site. The sphere radius was set to the reference length of the corresponding myosin or CL element, 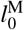 or 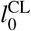, respectively. One candidate was selected randomly, and the element was attached to the two distinct AFs at its reference length. If no valid candidate was available, the element remained unbound.

#### 2.1.3. AF turnover and repositioning

AF turnover was modeled at a coarse-grained level through filament disassembly and repositioning, without explicitly representing molecular-scale polymerization and depolymerization (Maxian et al., 2021). Two AF-disassembly mechanisms were considered: lifetime-dependent disassembly modeled as a constant-rate stochastic process (Fig. 1(F)) and bending-dependent disassembly governed by the maximum bending angle of each AF (Fig. 1(G)). For AF *l*, the corresponding disassembly probabilities during a time step Δ*t* were given by

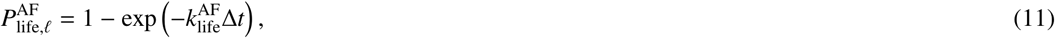

and

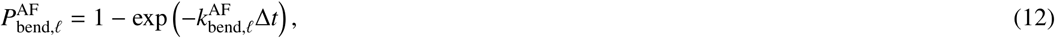

where 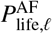 and 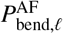 are the probabilities of lifetime-dependent and bending-dependent disassembly of AF *l*, respectively. Here, 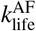 is the constant lifetime-dependent disassembly rate common to all AFs, whereas 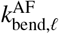 is the bending-dependent disassembly rate of AF *l*.

The bending-dependent disassembly rate of AF *l* was defined using a Hill function:

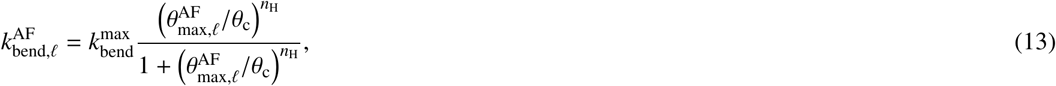

where 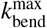 is the maximum bending-dependent disassembly rate, 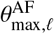 is the maximum angle between adjacent line elements within AF *l, θ*_c_ is the bending angle at which 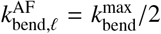, and *n*_H_ is the Hill coefficient. When an AF was disassembled, all CL and myosin elements attached to that AF were removed together with it. The bending-dependent disassembly rule introduced here provides a coarse-grained representation of the preferential removal of strongly bent or buckled AFs. This rule was motivated by experiments showing that myosin-induced buckling promotes severing and fragmentation at highly curved regions of AFs (Murrell and Gardel, 2012; Vogel et al., 2013), as well as by a previous model in which angle-dependent AF severing facilitated mechanical relaxation and cluster disassembly within contractile actomyosin clusters (Yu et al., 2018).

Spatial heterogeneity in cortical F-actin can influence the distribution of Rho activity and actin assembly through activator–inhibitor feedback (Bement et al., 2015; Maxian et al., 2025). Based on this biological motivation, we adopted a coarse-grained phenomenological rule in which disassembled AFs were preferentially repositioned to regions with low AF-node density. To evaluate the local AF-node density, an auxiliary triangular mesh was introduced on the spherical cortical surface. Each triangle is hereafter referred to as a surface mesh face, and multiple neighboring mesh faces were grouped into surface patches for spatial coarse-graining. For the *k*-th surface patch, the local AF-node density was defined as

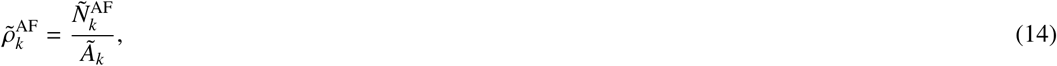

where 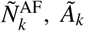, and 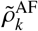 are the number of AF nodes, surface area, and AF-node density of the *k*-th surface patch, respectively. Surface patches within the lowest *q*_*ρ*_ quantile of the local AF-node-density distribution were selected as candidates for AF repositioning (Fig. 1(H)). A target mesh face was then selected randomly from the candidate patches with a probability proportional to its area. The first node of the repositioned AF was placed within the selected mesh face. The remaining nodes were arranged as a straight chain in a randomly selected direction, with adjacent nodes separated by the reference element length 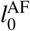 . If any node lay outside the spherical cortical region, a new direction was sampled until the entire AF was contained within the region.

To keep the number of AFs in the cortex constant, we prescribed the immediate repositioning of each disassembled AF within the cortical region using the procedure described above. Upon AF disassembly, all CL and myosin elements attached to the disassembled AF were transferred to their respective unbound pools. The subsequent recruitment of these elements to the cortical network was governed by their respective stochastic binding processes. Thus, the model conserved the total numbers of CL and myosin elements, including both cortex-bound and pooled elements, while allowing the numbers of cortex-bound elements to vary over time.

The membrane constraints representing ERM-mediated anchoring that were lost upon AF disassembly were reassigned immediately after AF repositioning. Specifically, a number of distinct AF nodes equal to the number of lost constraints were selected uniformly at random from unconstrained AF nodes near the fixed spherical boundary. Tangential sliding constraints were assigned to the selected nodes within the same turnover event.

### 2.2. Evaluation of cortical tension at the cellular scale

To evaluate the cortical tension generated by the actomyosin network, the local surface stress tensor was reconstructed on each surface mesh face. The same auxiliary triangular mesh introduced for evaluating the local AF-node density was used for this analysis. For each mesh edge *s*, a local section plane containing the edge was defined, and the line elements representing AFs, CLs, and myosin minifilaments were projected onto this plane. The intersections of the projected line elements with the finite-length mesh edge were then identified.

The internal force transmitted across edge *s* was expressed as

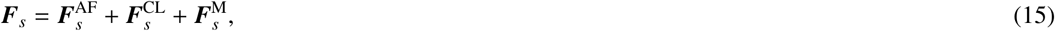

where 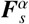 is the resultant internal force transmitted across edge *s* by component type *α* ∈ {AF, CL, M}. Each contribution was obtained by summing the forces transmitted by the corresponding elements intersecting the edge, using the force definition in Eq. (5). For each face *f* adjacent to edge *s*, the internal force ***F***_*s*_ was divided by the edge length *L*_*s*_ and projected onto the local tangent plane of the face to obtain the in-plane traction vector 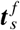 . The outward unit conormal to edge *s* in the tangent plane of face *f* was denoted by 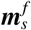.

The local surface stress tensor τ _*f*_ was reconstructed from the traction vectors on the three edges of face *f* . Assuming a symmetric surface stress tensor, τ _*f*_ was obtained by solving the following least-squares problem:

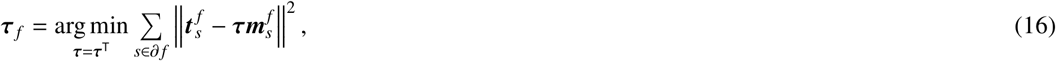

where *∂ f* denotes the set of the three edges of face *f* . The three edge tractions provide six in-plane traction components for determining the three independent components of the symmetric surface stress tensor.

The local cortical tension *γ*_*f*_ on face *f* was defined as the isotropic component of the surface stress tensor:

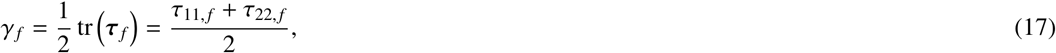

where *τ*_11, *f*_ and *τ*_22, *f*_ are the two normal components of τ _*f*_ in the local tangent basis of face *f* .

The cellular-scale cortical tension was defined as the area-weighted mean of the local cortical tension over the spherical cortical surface:

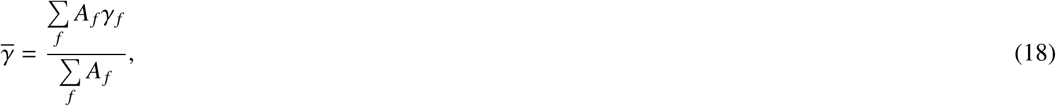

where *A*_*f*_ is the area of surface mesh face *f* .

### 2.3. Evaluation of component distributions, tensile energy, and network connectivity

To quantify the spatial heterogeneity of the AF distribution, AF nodes were assigned to the nearest surface mesh face. The AF-node density on face *f* was defined as

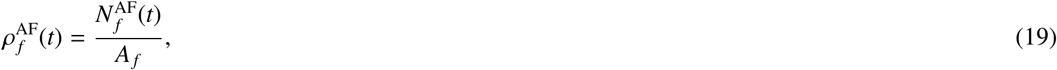

where 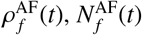, and *A*_*f*_ are the AF-node density, number of AF nodes assigned to face *f*, and area of face *f* at time *t*, respectively. The area-weighted spatial mean of the AF-node density was calculated as

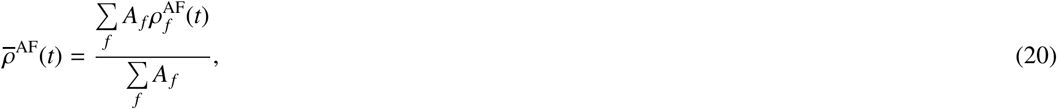

and its spatial coefficient of variation was defined as

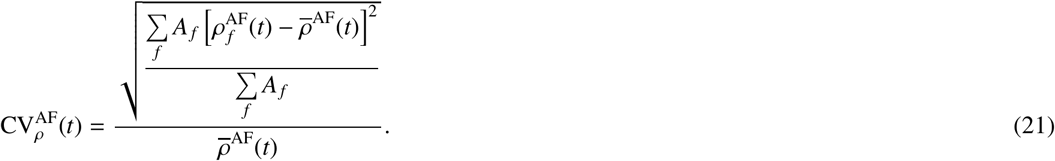

This coefficient quantifies the spatial heterogeneity of the AF-node density, with larger values indicating a more hetero-geneous AF distribution.

To quantify network connectivity, we constructed an undirected graph in which each AF was represented as a graph node and connections between distinct AFs mediated by bound CL or myosin elements were represented as graph edges.

Denoting the set of AFs belonging to the largest connected component by *V*_LCC_(*t*), the LCC fraction was defined as

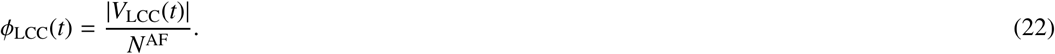

The LCC fraction represents the fraction of all AFs contained in the largest connected network and was used to quantify the global connectivity of the cortical AF network.

To quantify the relative spatial distribution and mechanical loading of each component, we calculated the element fraction and tensile energy per element on each surface mesh face. These quantities were evaluated separately for component type *α* ∈ {AF, CL, M}, considering only bound CL and myosin elements. For AFs, each line element was treated as an individual element. The portion of each element assigned to a surface mesh face was determined using the same local projection procedure as that used in the surface-stress calculation.

For element *e* of component type *α*, the weight assigned to face *f* was defined as

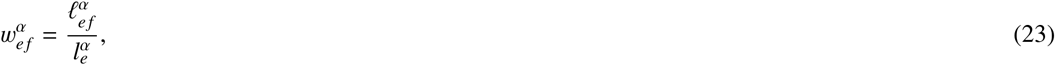

where 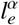 is the total length of element *e* and 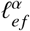 is the length of its portion assigned to face *f* . The equivalent element count on face *f* was defined as

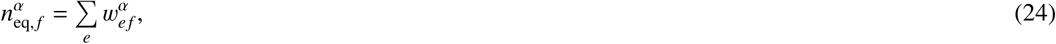

where the sum was taken over all elements of component type *α*.

The element fraction of component type *α* on face *f* was defined as

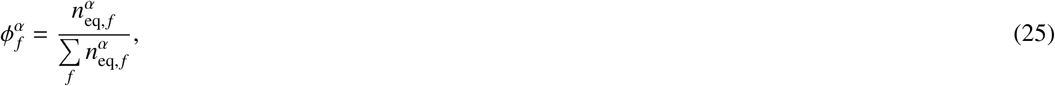

where the sum over *f* includes all surface mesh faces. Thus, 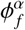 represents the fraction of the total element content of component type *α* assigned to face *f* .

The stretching energy of each element was allocated to the surface mesh faces using the same weights. The tensile energy per element of component type *α* on face *f* was defined as

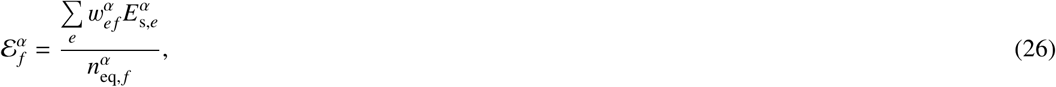

where 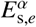 is the stretching energy of element *e* of component type *α*, as defined in Eq. (2). The tensile energy per element was left undefined on faces with 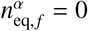 .

### 2.4. Simulation setup

In the initial configuration, each AF was placed as a straight chain whose line elements were set to their reference length 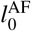 . The first node of each AF was placed uniformly at random within the spherical cortical region, and the direction of the AF was selected randomly. If any node lay outside the cortical region, a new direction was sampled until the entire AF was contained within the region. The polarity of each AF was defined according to the ordering of its nodes.

After all AFs had been placed, CL and myosin elements were bound to geometrically accessible pairs of distinct AFs using the binding procedure described above. Their initial lengths were set to their respective reference lengths, 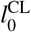 and 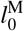. To represent ERM-mediated membrane anchoring, a prescribed number of AF nodes, *N*^ERM^ = 10000, near the fixed spherical boundary were selected uniformly at random for tangential sliding constraints.

The model and analysis parameters are listed in Table 1. Each simulation was performed for a total duration of *T*_sim_ = 200 s.

**Table 1.**
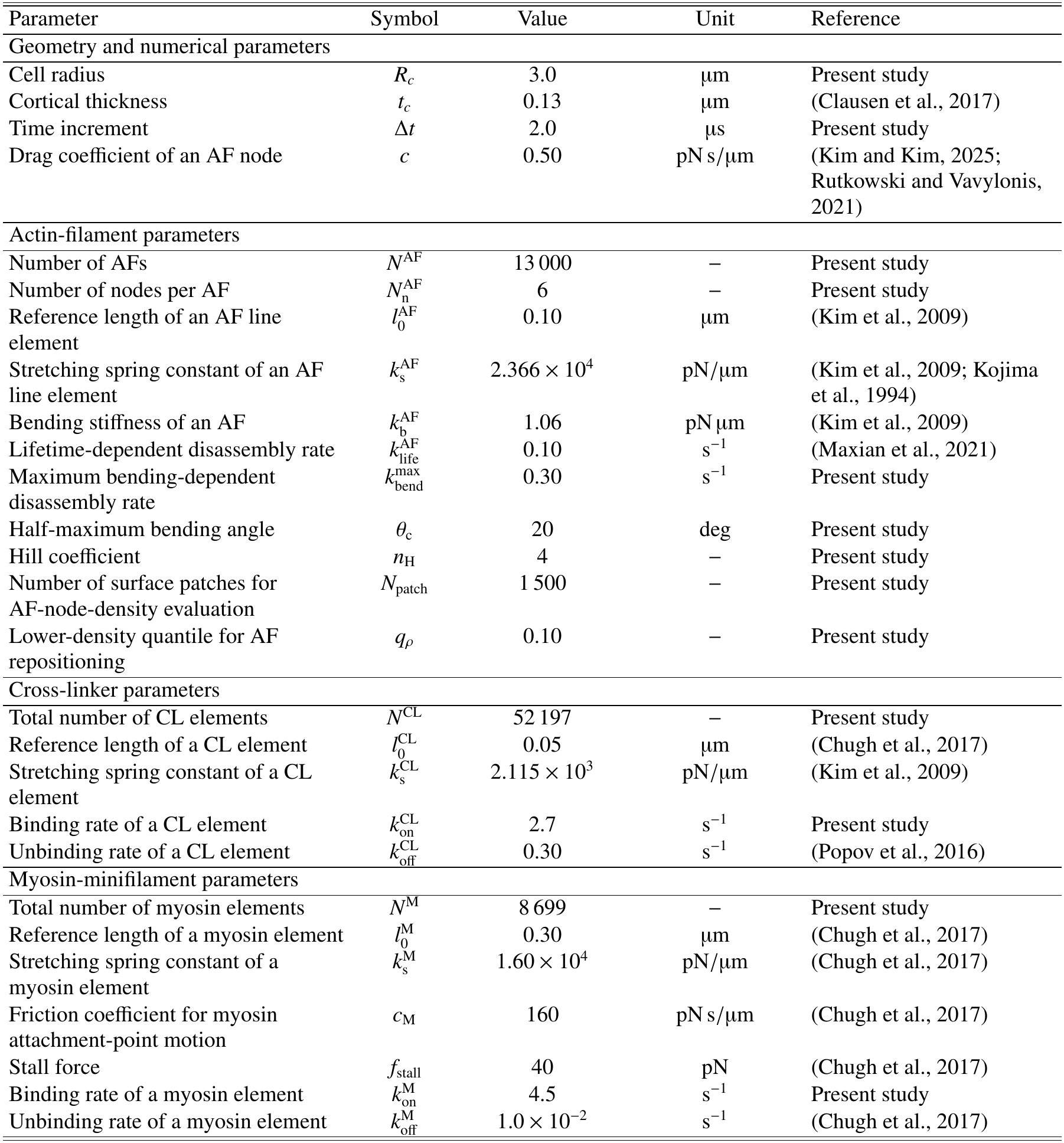
Model and analysis parameters.

| Parameter | Symbol | Value | Unit | Reference |
|---|---|---|---|---|
| Geometry and numerical parameters |  |  |  |  |
| Cell radius | *R*<sub>*c*</sub> | 3.0 | µm | Present study |
| Cortical thickness | *t*<sub>*c*</sub> | 0.13 | µm | (Clausen et al., 2017) |
| Time increment | Δ*t* | 2.0 | µs | Present study |
| Drag coefficient of an AF node | *c* | 0.50 | pN s/µm | (Kim and Kim, 2025; Rutkowski and Vavylonis, 2021) |
| Actin-filament parameters |  |  |  |  |
| Number of AFs | *N*<sup>AF</sup> | 13 000 | – | Present study |
| Number of nodes per AF | *N*<sub>n</sub><sup>AF</sup> | 6 | – | Present study |
| Reference length of an AF line element | *l*<sub>0</sub><sup>AF</sup> | 0.10 | µm | (Kim et al., 2009) |
| Stretching spring constant of an AF line element | *k*<sub>s</sub><sup>AF</sup> | 2.366 × 10<sup>4</sup> | pN/µm | (Kim et al., 2009; Kojima et al., 1994) |
| Bending stiffness of an AF | *k*<sub>b</sub><sup>AF</sup> | 1.06 | pN µm | (Kim et al., 2009) |
| Lifetime-dependent disassembly rate | *k*<sub>life</sub><sup>AF</sup> | 0.10 | s<sup>−1</sup> | (Maxian et al., 2021) |
| Maximum bending-dependent disassembly rate | *k*<sub>bend</sub><sup>max</sup> | 0.30 | s<sup>−1</sup> | Present study |
| Half-maximum bending angle | *θ*<sub>c</sub> | 20 | deg | Present study |
| Hill coefficient | *n*<sub>H</sub> | 4 | – | Present study |
| Number of surface patches for AF-node-density evaluation | *N*<sub>patch</sub> | 1 500 | – | Present study |
| Lower-density quantile for AF repositioning | *q*<sub>*ρ*</sub> | 0.10 | – | Present study |
| Cross-linker parameters |  |  |  |  |
| Total number of CL elements | *N*<sup>CL</sup> | 52 197 | – | Present study |
| Reference length of a CL element | *l*<sub>0</sub><sup>CL</sup> | 0.05 | µm | (Chugh et al., 2017) |
| Stretching spring constant of a CL element | *k*<sub>s</sub><sup>CL</sup> | 2.115 × 10<sup>3</sup> | pN/µm | (Kim et al., 2009) |
| Binding rate of a CL element | *k*<sub>on</sub><sup>CL</sup> | 2.7 | s<sup>−1</sup> | Present study |
| Unbinding rate of a CL element | *k*<sub>off</sub><sup>CL</sup> | 0.30 | s<sup>−1</sup> | (Popov et al., 2016) |
| Myosin-minifilament parameters |  |  |  |  |
| Total number of myosin elements | *N*<sup>M</sup> | 8 699 | – | Present study |
| Reference length of a myosin element | *l*<sub>0</sub><sup>M</sup> | 0.30 | µm | (Chugh et al., 2017) |
| Stretching spring constant of a myosin element | *k*<sub>s</sub><sup>M</sup> | 1.60 × 10<sup>4</sup> | pN/µm | (Chugh et al., 2017) |
| Friction coefficient for myosin attachment-point motion | *c*<sub>M</sub> | 160 | pN s/µm | (Chugh et al., 2017) |
| Stall force | *f*<sub>stall</sub> | 40 | pN | (Chugh et al., 2017) |
| Binding rate of a myosin element | *k*<sub>on</sub><sup>M</sup> | 4.5 | s<sup>−1</sup> | Present study |
| Unbinding rate of a myosin element | *k*<sub>off</sub><sup>M</sup> | 1.0 × 10<sup>−2</sup> | s<sup>−1</sup> | (Chugh et al., 2017) |

## 3. Results

### 3.1. Spatial organization and dynamics of cortical tension

Local contractile forces generated by myosin minifilaments stretched the AFs and CLs, resulting in the formation of a tension-bearing network throughout the cell cortex (Fig. 2(A) and (B)). The local cortical tension was spatially heterogeneous, and a network-like distribution of high-tension regions developed over time. The spatial distribution of the number of AF nodes assigned to each surface mesh face 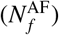 did not obviously exhibit the network-like organization observed in the tension distribution (Fig. 2(C)). To quantify the connectivity, global mechanical state, and spatial heterogeneity of the cortical network, we evaluated the temporal evolutions of the fraction of AFs belonging to the largest connected component (*ϕ*_LCC_), the area-weighted spatial average of the local cortical tension 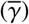, and the spatial coefficient of variation of the AF-node density 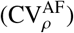 (Fig. 2(D)–(F)). *ϕ*_LCC_ decreased from nearly unity at the initial state to approximately 0.96 within the first second and subsequently remained near this value, indicating that a globally connected network containing most of the AFs was maintained (Fig. 2(D)). After the initial transient period, 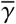 and 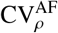 also remained nearly constant over time (Fig. 2(E) and (F)).

**Fig. 2.**
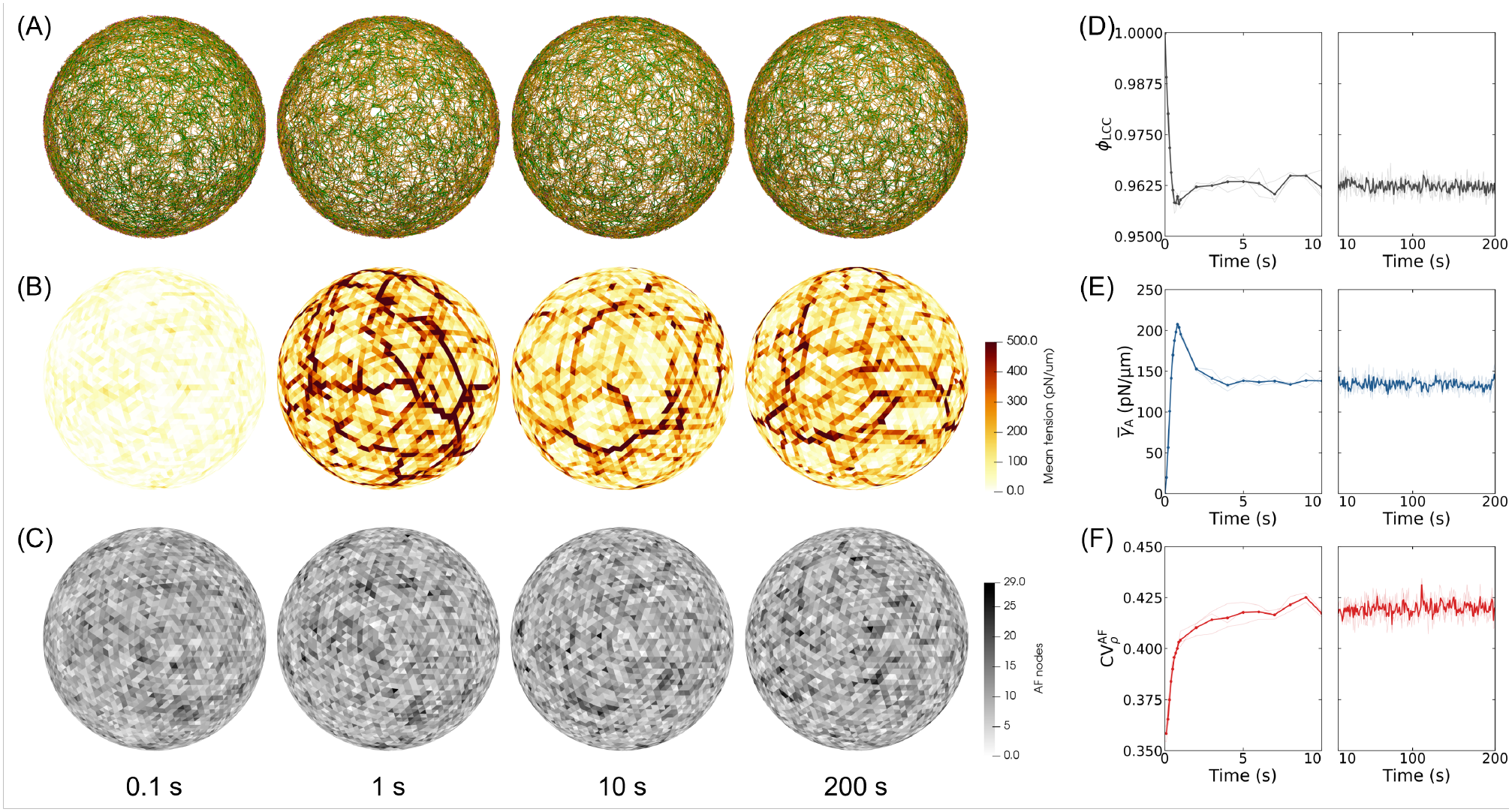
Spatial organization and dynamics of the actomyosin cortex. (A) Actomyosin network structure. (B) Local cortical tension distribution. (C) Number of AF nodes assigned to each mesh face 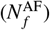 . (D)–(F) Time courses of (D) the fraction of AFs in the largest connected component (*ϕ*_LCC_), (E) the area-weighted mean cortical tension 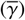, and (F) the spatial coefficient of variation of AF-node density 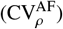. In (D)–(F), left and right plots show 0–10 s and 10–200 s, respectively; thin lines show three independent simulations and thick lines show their mean.

To identify the local structural and mechanical quantities associated with cortical tension (*γ*_*f*_ ), we evaluated the element fraction 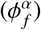 and tensile energy per element 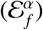 on each surface mesh face at 200 s (Figs. 3(A) and (B)). These distributions were compared with the local mean cortical tension at the same time point shown in Fig. 2(B). We then calculated area-weighted Spearman rank correlations between local mean cortical tension and these two quantities (Fig. 3(C)). For direct comparison among AFs, CLs, and myosin, all correlations were evaluated over the same set of mesh faces with positive local mean cortical tension on which all three components were present. The correlations between element fraction and local mean cortical tension were 0.251 ± 0.015 for AFs, 0.298 ± 0.010 for CLs, and 0.254 ± 0.022 for myosin. By contrast, the correlations with tensile energy per element were 0.560 ± 0.009 for AFs, 0.490 ± 0.007 for CLs, and 0.505 ± 0.017 for myosin. For all three components, tensile energy per element showed a stronger positive correlation with local mean cortical tension than did element fraction, with the strongest correlation observed for AFs.

**Fig. 3.**
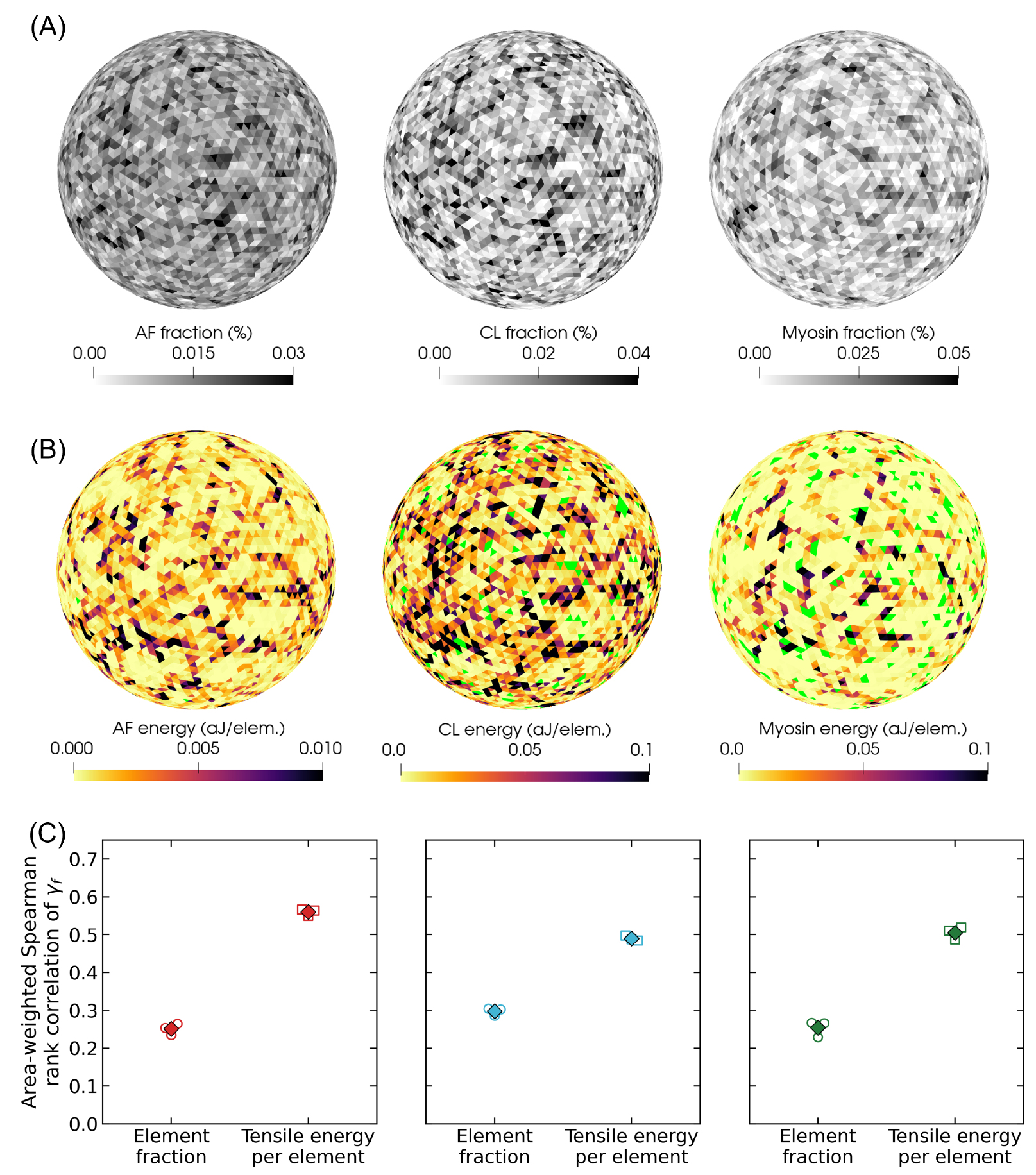
Spatial relationships between cortical tension, component abundance, and tensile energy at 200 s. (A) Element fraction 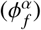 and (B) tensile energy per element 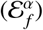 for AFs, bound CLs, and bound myosin, shown from left to right. Light-green regions in (B) contain no assigned elements, for which ^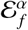^ is undefined. (C) Area-weighted Spearman rank correlations of local mean cortical tension (*γ*_*f*_ ) with 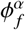 and ^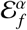^ for AFs, CLs, and myosin, shown from left to right. All correlations were evaluated over the same set of mesh faces with *γ*_*f*_ *>* 0 and all three components present. Open symbols indicate individual simulations, and diamonds indicate their means.

We focused on high-stretch AF elements and examined their spatial occurrence and orientation relative to the first-principal-tension orientation of τ _*f*_ (Fig. 4). The surface-area fraction of mesh faces containing high-stretch AF elements increased with first principal tension, from approximately 10.4% in the lowest-tension decile to 87.2% in the highest-tension decile (Fig. 4(B)). The mean angular difference between the high-stretch-AF director and the first-principal-tension orientation decreased from the low-tension range and remained at approximately 27–29^°^ over the intermediate- to high-tension range (Fig. 4(C)). In the highest-tension decile, the mean angular difference was approximately 27.6^°^ for high-stretch AF elements and 36.4^°^ for all AF elements on the same mesh faces. Accordingly, high-stretch AF elements were more strongly aligned with the first-principal-tension orientation than the local AF population as a whole.

**Fig. 4.**
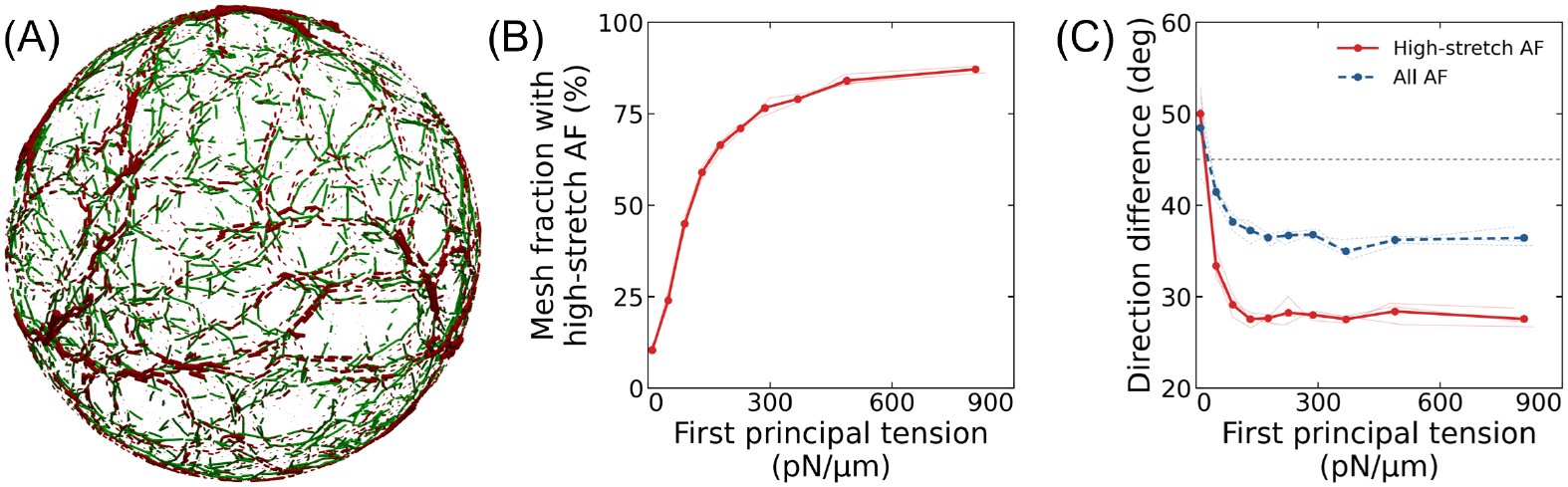
Local cortical tension and AF orientation at 200 s. (A) High-stretch AF elements (green), defined as the top 10% by stretch ratio in each simulation, and first-principal-tension orientations (red line segments), with line lengths scaled by the magnitude of the first principal tension. (B) Surface-area fraction of mesh faces containing high-stretch AF elements in each first-principal-tension decile. (C) Area-weighted mean angular difference between the first-principal-tension orientation and local directors of all AF elements (blue) or high-stretch AF elements (red), evaluated on the same faces containing high-stretch AF elements. In (B) and (C), horizontal coordinates show the area-weighted mean first principal tension in each decile; thin lines show three independent simulations and thick lines with markers show their mean. The dashed line in (C) marks 45^°^, the expectation for random in-plane orientations.

We next examined how the local cortical tension pattern and cortical components changed over time while the cell-scale cortical tension remained approximately constant. The analysis focused on the quasi-steady-state interval from 180 to 200 s. We quantified the temporal autocorrelation of the local cortical tension field by calculating the area-weighted Pearson correlation across mesh faces between the tension distributions at times *t* and *t* + Δ*t*. The persistence fraction was defined as the fraction of AFs present at time *t* that underwent no disassembly, or of CLs and myosin minifilaments bound at time *t* that underwent no detachment, during the interval from *t* to *t* + Δ*t*. For each time lag Δ*t*, both the correlations and persistence fractions were averaged over time origins spaced Δ*t* apart, giving non-overlapping intervals within the analysis period. For each quantity, the characteristic time *τ*_1*/e*_ was defined as the time lag at which the correlation or persistence fraction decreased to 1*/e*.

Although the cell-scale cortical tension remained approximately constant during the quasi-steady state, the temporal autocorrelation of the local cortical tension field decreased rapidly with increasing time lag, yielding a characteristic time of 0.392 ± 0.005 s (Fig. 5(A) and (C)). Consistent with this correlation-based characteristic time, direct comparison of the tension maps showed that the spatial tension pattern was substantially reorganized over a comparable subsecond timescale (Fig. 5(D)). The cortical components exhibited distinct persistence times: 4.748 ± 0.033 s for AFs, 1.331 ± 0.008 s for CLs, and 0.488 ± 0.001 s for myosin (Fig. 5(B) and (C)). Thus, the local tension pattern was reorganized substantially faster than AFs and CLs, whereas its characteristic time was closest to the persistence time of myosin.

**Fig. 5.**
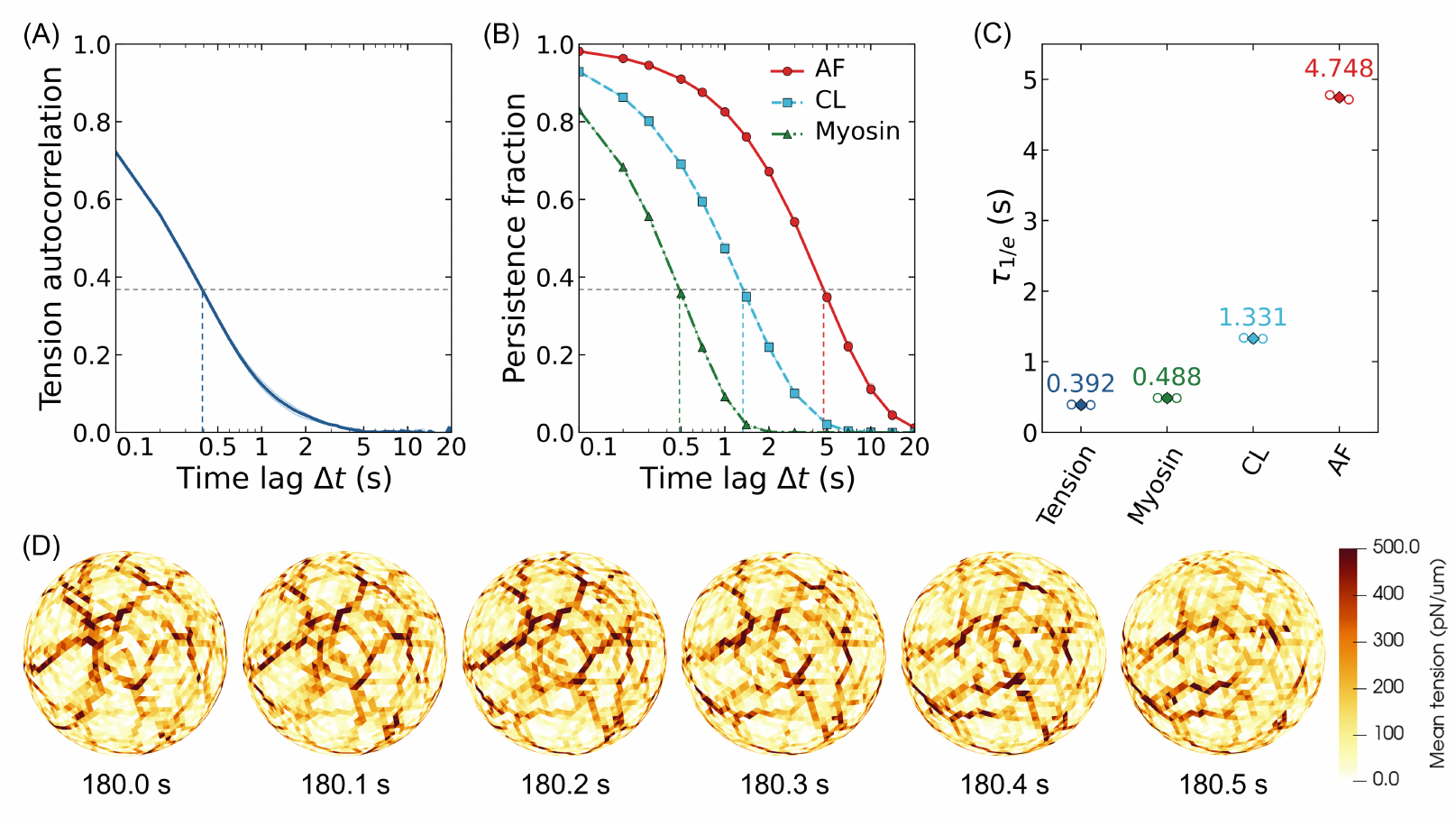
Timescales of cortical tension reorganization and component persistence. Analyses used the quasi-steady-state interval of 180–200 s at a temporal resolution of 0.1 s. (A) Temporal autocorrelation of local cortical tension. (B) Persistence fractions of AFs, CLs, and myosin. (C) Characteristic times *τ*_1*/e*_ at which the autocorrelation or persistence fraction decreased to 1*/e*. (D) Short-time changes in the local cortical tension pattern. In (A) and (B), thin lines show three independent simulations and thick lines show their mean; horizontal and vertical dashed lines indicate 1*/e* and the corresponding *τ*_1*/e*_ from the mean curves, respectively. In (C), open circles show individual simulations and diamonds show their mean.

### 3.2. Effect of AF turnover on tension maintenance

To assess the roles of the AF-relocation rule and bending-dependent disassembly, we compared conditions in which either mechanism was altered. The reference condition combined preferential AF relocation to regions of low AF density with both lifetime-dependent and bending-dependent disassembly. Two comparison conditions were considered: a uniform-relocation condition, in which disassembled AFs were repositioned uniformly at random, and a lifetime-only condition, in which preferential relocation to low-density regions was retained but bending-dependent disassembly was omitted.

For uniform relocation, the first node of each repositioned AF was placed uniformly at random within the spherical cortical region, independently of the local AF-node density. For the lifetime-only condition, the overall AF-disassembly frequency was adjusted to be comparable to that under the reference condition, allowing the effect of bending-dependent disassembly to be examined without a substantial change in the total disassembly frequency. For this adjustment, the effective AF-disassembly rate was estimated from the reference-condition simulations by dividing the number of AF-disassembly events by the cumulative time over which the AFs were present in the cortex. The lifetime-dependent disassembly rate was then set to the estimated effective rate, 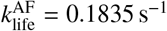.

Under the uniform relocation condition, the network snapshots showed more pronounced local AF concentration and a more heterogeneous density distribution than under the reference condition (Fig. 6(A)). Local AF concentration was also observed under the lifetime-only condition, although it was less pronounced than under the uniform relocation condition. A network-like distribution of high-tension regions formed under all conditions; however, the branching density of the high-tension network and the magnitude of the local cortical tension differed among the conditions (Fig. 6(B)).

**Fig. 6.**
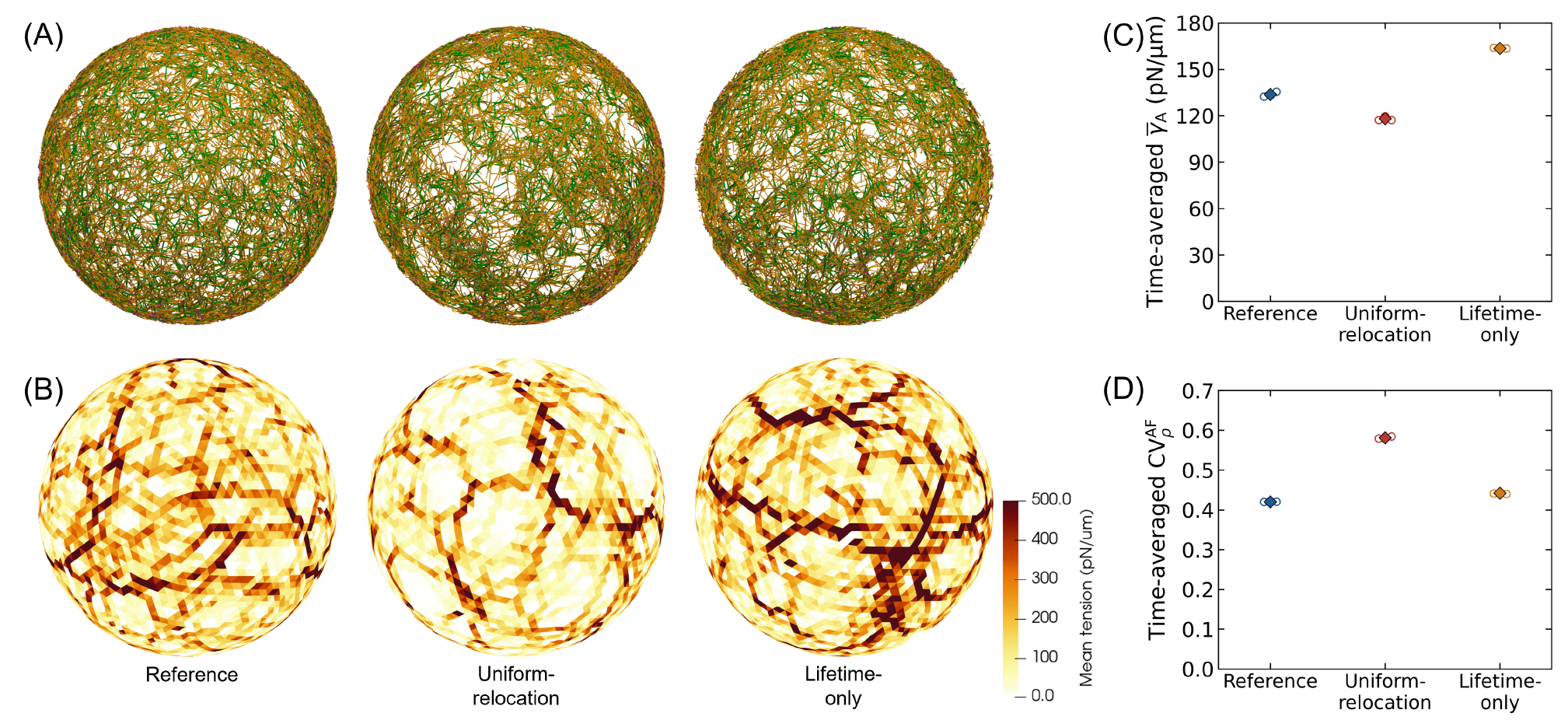
Effects of AF relocation and bending-dependent disassembly on cortical tension. (A) Actomyosin network structure and (B) local cortical tension distribution under the reference (left), uniform-relocation (center), and lifetime-only (right) conditions. (C) Spatiotemporal mean cortical tension and (D) time-averaged spatial coefficient of variation of AF-node density, evaluated over 101–200 s. Open circles show individual simulations; diamonds and error bars show the mean and standard deviation of three independent simulations.

For quantitative comparison, the area-weighted spatial average of the local cortical tension 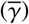 and the spatial co-efficient of variation of the AF-node density 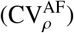 were calculated at each time point and subsequently averaged over 101–200 s for each simulation. The resulting time-averaged values were compared among the three turnover conditions using three independent simulations for each condition. The spatiotemporally averaged cortical tension was lower under the uniform-relocation condition and higher under the lifetime-only condition, relative to the reference condition (Fig. 6(C)). AF-node density variation was greatest under the uniform-relocation condition, followed by the lifetime-only and reference conditions (Fig. 6(D)).

### 3.3. Effects of equatorial myosin localization and AF accumulation

To examine how different modes of equatorial cortex organization affect network mechanics and dynamics, we compared two idealized conditions: direct myosin localization and AF accumulation. In both conditions, the localization region was defined as an equatorial band spanning absolute latitudes below 10^°^.

Under the direct myosin-localization condition, AF line elements within the equatorial band were assigned a fourfold higher relative weight than those outside the band when selecting the first attachment site of a myosin minifilament, without directly altering the AF distribution. This modification was applied only to myosin minifilaments and not to CLs. The procedure for selecting the second attachment site and the mechanical properties of bound myosin elements were unchanged from those under the reference condition. This condition therefore represented an idealized case in which force-generating elements preferentially localized on the existing cortical AF network.

Under the AF-accumulation condition, a probability parameter *α* was introduced to control the fraction of disassembled AFs preferentially relocated to the equatorial band. After the onset of AF accumulation, each disassembled AF was assigned to equatorial relocation with probability *α*; otherwise, it was repositioned according to the low-density relocation rule used under the reference condition. For equatorial relocation, the first node was placed within the equatorial band, and the remaining nodes were positioned using the same AF-placement procedure described above. The value *α* = 0.40 was chosen so that the AF-accumulation condition yielded approximately the same number of bound myosin minifilaments within the equatorial band as the direct myosin-localization condition.

An equatorward cortical flow developed under both conditions, as evaluated from tangential AF velocities spatially averaged within each mesh face and temporally averaged over 120–200 s (Fig. 7(A)). For quantitative comparisons, the equatorward components of the mesh-face-averaged velocities were further averaged over the region of interest using mesh-face areas as weights. Latitudinal profiles were obtained by averaging the equatorward velocity components within each absolute-latitude bin. Under both localization conditions, the equatorward velocity was higher near the equator and decreased toward the poles, whereas the no-imposed-localization condition exhibited little net flow (Fig. 7(B)). For comparison with experimental measurements (Zhou and Wang, 2008; Reymann et al., 2016), the averaging region was instead defined as the absolute-latitude range from 10^°^ to 45^°^ in both hemispheres. The simulated equatorward AF velocities were of the same order of magnitude as the experimental values (Fig. 7(C)).

**Fig. 7.**
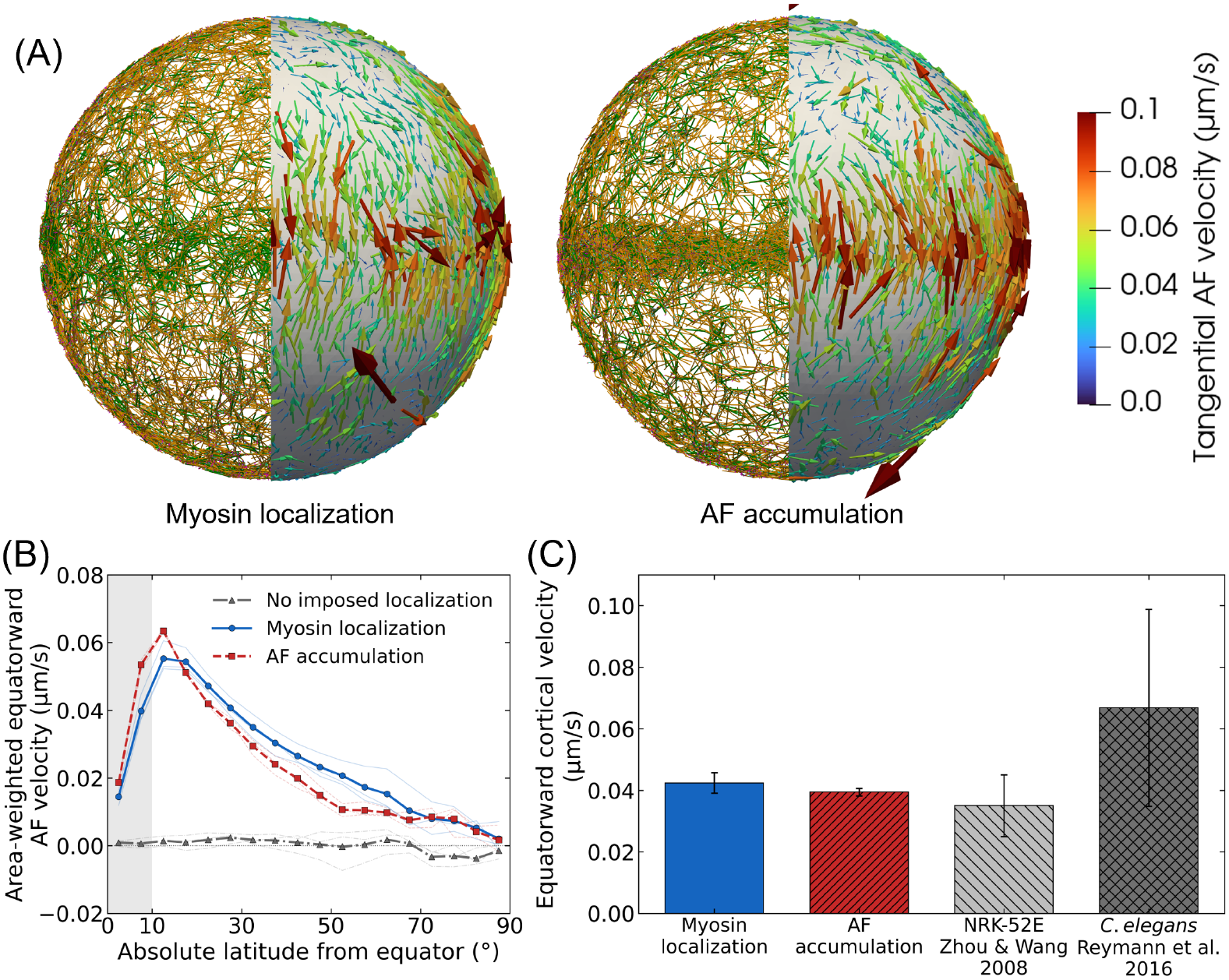
Cortical flow under equatorial localization. (A) Cortical network structure and tangential AF velocity field. Left and right panels show the direct myosin-localization and AF-accumulation conditions, respectively. (B) Latitudinal profiles of equatorward AF velocity under both localization conditions and the no-imposed-localization condition. Gray shading indicates the localization region at absolute latitudes below 10^°^. Thin lines show three independent simulations; thick lines with markers show their mean. (C) Equatorward AF velocities averaged over absolute latitudes from 10^°^ to 45^°^ in both hemispheres, compared with experimental measurements (gray bars) (Zhou and Wang, 2008; Reymann et al., 2016). Velocities were averaged over 120–200 s; spatial averages in (B) and (C) were area-weighted. Error bars indicate the standard deviation among simulations or the reported experimental variability.

Although equatorward cortical flow developed under both conditions, the spatial organization of high-tension regions near the equatorial band differed between the two localization mechanisms (Fig. 8(A)). Under the direct myosin-localization condition, the branching density of the high-tension network increased within the equatorial band. In contrast, under the AF-accumulation condition, continuous high-tension structures aligned with the equatorial band were formed. To compare the latitudinal distributions of cortical tension and cortical flow, we evaluated the meridional component, *τ*_*θθ*_, and the circumferential component, *τ*_*ϕϕ*_, of the cortical stress tensor as functions of absolute latitude (Fig. 8(B,C)). Under the no-imposed-localization condition, both components were approximately uniform with latitude and had similar values. Under both localization conditions, *τ*_*θθ*_ decreased from the equator toward the poles, but the profiles differed. Under direct myosin localization, it decreased gradually over a broad range of latitudes, whereas under AF accumulation, it decreased sharply near the equatorial band and was relatively flat at higher latitudes. In both conditions, *τ*_*ϕϕ*_ was elevated within the equatorial band, with a more pronounced increase under AF accumulation. Outside the band, *τ*_*ϕϕ*_ was relatively uniform with latitude, and *τ*_*θθ*_ exceeded *τ*_*ϕϕ*_ under both localization conditions.

**Fig. 8.**
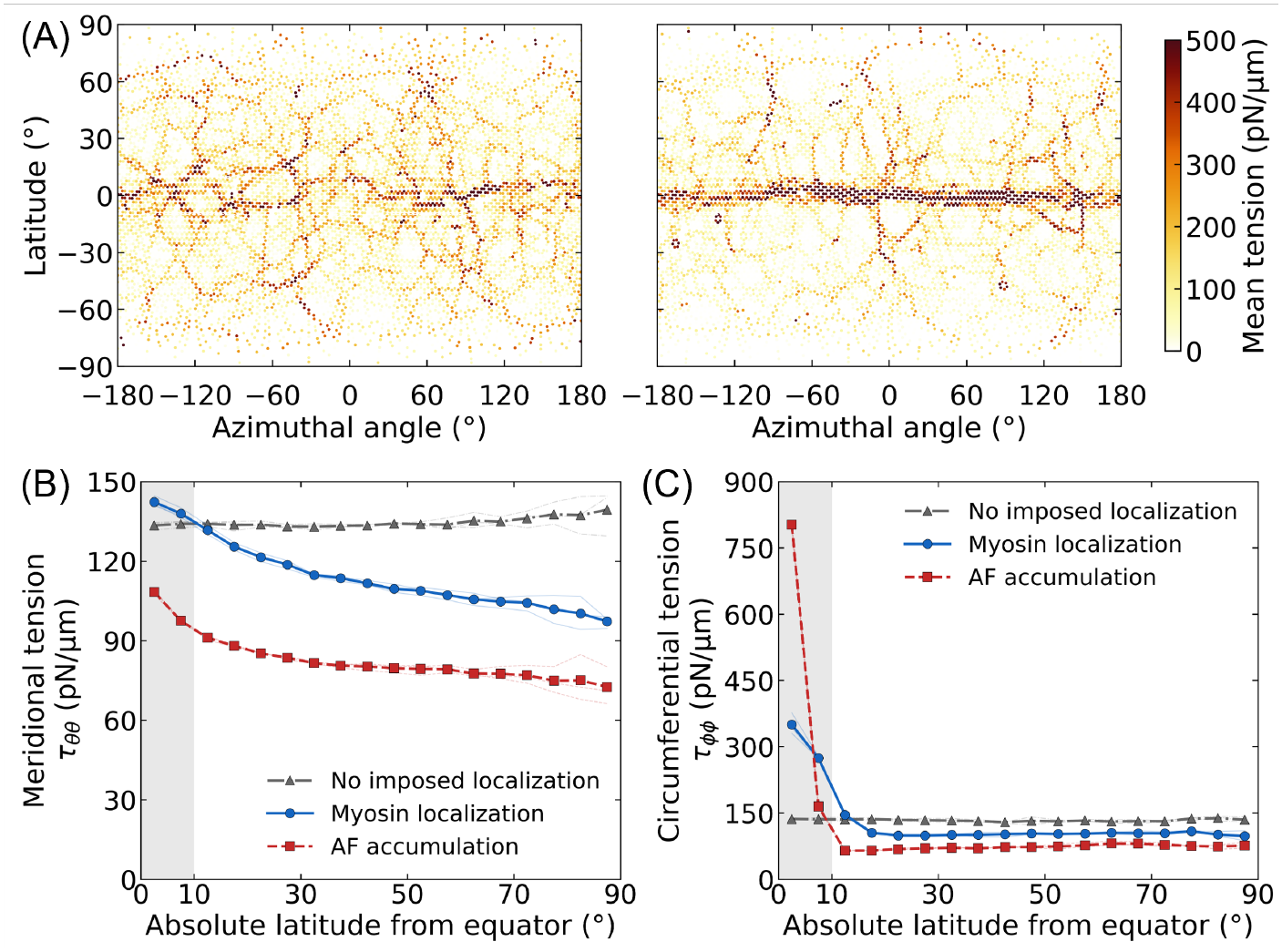
Cortical tension organization under equatorial localization. (A) Local mean cortical tension at 200 s. Left and right panels show the direct myosin-localization and AF-accumulation conditions, respectively. (B) Meridional tension *τ*_*θθ*_ and (C) circumferential tension *τ*_*ϕϕ*_ as functions of absolute latitude under both localization conditions and the no-imposed-localization condition. Tensions in (B) and (C) were averaged over 120–200 s and area-weighted within each absolute-latitude bin. Gray shading indicates the localization region at absolute latitudes below 10^°^. Thin lines show three independent simulations; thick lines with markers show their mean.

## 4. Discussion

The local AF element fraction was only weakly correlated with local mean cortical tension, whereas tensile energy per element showed stronger positive correlations with local mean cortical tension than did element fraction for all three cortical components, with the strongest correlation observed for AFs (Fig. 3). Consistent with this pattern, high-stretch AF elements occurred preferentially in high-tension regions and showed greater alignment with the first-principal-tension orientation (Fig. 4). These results indicate that the spatial organization of cortical tension is associated not only with local component abundance but also with filament-scale mechanical states, particularly AF stretch and orientation. In laser-ablation experiments, local cutting of the prestressed cortex releases stored stress, causing the cut edges to recoil. Differences in initial recoil velocity with cutting direction have been reported to reflect the anisotropy of cortical tension before ablation (Mayer et al., 2010; Spira et al., 2017). FRET-based tension-sensor studies have also shown that molecular loading in cortical actin and stress fibers responds differently depending on the direction of external stretch (Amiri et al., 2023). Taken together, these experimental findings and our results suggest that loaded, oriented AFs transmit force anisotropically, contributing to the network-like distribution of high-tension regions.

The load-bearing pathways were not maintained as fixed assemblies of components that continuously persisted or remained bound. Over a 20-s time lag, the fraction of AFs that had not disassembled and the fractions of CLs and myosin minifilaments that were bound at time *t* and had not detached both decreased to nearly zero. In contrast, the fraction of AFs in the largest connected component remained approximately 0.96, while cell-scale cortical tension and the spatial coefficient of variation of AF-node density remained nearly constant after the initial transient period (Figs. 2(D–F) and 5(B)). The decay of the temporal correlation of the local tension pattern suggests that high-tension pathways changed while global connectivity, AF-density heterogeneity, and cell-scale cortical tension remained stable. A previous model showed that, under suitable turnover conditions, filament turnover can relax elastic resistance without substantial loss of network connectivity and sustain active stress (McFadden et al., 2017). Although the turnover mechanism in that model differs from the AF disassembly and repositioning used here, its findings are functionally consistent with our interpretation that turnover can contribute to maintaining the mechanical function of the network.

The timescale analysis further provides insight into the dynamics underlying this reorganization. The characteristic time of the tension pattern was much shorter than the persistence times of AFs and CLs but was closest to that of myosin (Fig. 5(C)). Because myosin binding, unbinding, and movement directly change where contractile forces act within the AF network, this temporal correspondence suggests that myosin dynamics may contribute to the rapid redistribution of force and reorganization of the local tension pattern. This interpretation is mechanically consistent with experimental findings that changes in cortical myosin recruitment, activity, and overlap with the AF network alter cortical tension (Ramanathan et al., 2015; Truong Quang et al., 2021), and with analyses linking spatial variations in myosin-dependent contractility to cortical tension and flow (Mayer et al., 2010). However, the similarity of the characteristic times alone does not establish a causal relationship; direct perturbation of myosin dynamics would be required to test this interpretation. Taken together, the present results suggest a nested dynamical organization in which myosin dynamics rapidly reorganize where forces are transmitted within the AF network, while the slower AF disassembly and repositioning may contribute to renewing the network that maintains global connectivity and cell-scale cortical tension.

The turnover conditions differed in both AF-density heterogeneity and cell-scale cortical tension. Under the uniform-relocation condition, local AF concentration was more pronounced and the spatial coefficient of variation of AF-node density was the highest among the three conditions, whereas cell-scale cortical tension decreased (Fig. 6). In the reference condition, our model incorporated a phenomenological rule that preferentially repositioned AFs to low-density regions, inspired by a negative-feedback circuit in which active Rho promotes F-actin assembly and the assembled F-actin suppresses Rho activity. By contrast, under the uniform-relocation condition, AFs were reintroduced independently of the existing AF-density distribution, so no mechanism acted to correct density biases generated during contraction. This may have allowed local AF concentration and spatial gaps to persist, potentially impairing load transmission. Previous models have likewise shown that insufficient force relaxation through AF turnover can produce heterogeneous networks with large AF clusters (Yu et al., 2018), and that the spatial rules for introducing new filaments affect bundle formation and sustained contractility (Tam et al., 2026). These results and previous modeling studies suggest that spatial feedback responsive to AF density may suppress density biases and contribute to maintaining load transmission and cell-scale cortical tension.

Under the lifetime-only condition, local AF concentration and AF-density heterogeneity increased slightly relative to the reference condition. In contrast, cell-scale cortical tension increased markedly, and the spatial contrast in the local tension distribution also became more pronounced (Fig. 6). Experiments have shown that myosin-induced AF buckling promotes severing and fragmentation at highly curved regions (Murrell and Gardel, 2012; Vogel et al., 2013). The bending-dependent disassembly rule in our model is a coarse-grained representation of the selective removal of strongly bent or buckled AFs. Because this disassembly mechanism was absent under the lifetime-only condition, strongly deformed AFs remained in the network until lifetime-dependent disassembly and may have continued to bear load while mechanically connected, thereby contributing to the increase in cortical tension. A previous model also showed that infrequent angle-dependent AF severing leads to the formation of large, long-lived clusters, qualitatively consistent with the increased local AF concentration observed here (Yu et al., 2018). These findings suggest that bending-dependent AF removal may help limit the accumulation of strongly deformed AFs and regulate local and cell-scale cortical tension.

The opposite changes in tension under the two comparison conditions suggest that the mechanical state of the cortex cannot be explained solely by AF concentration or density heterogeneity, but is also related to AF deformation and load-bearing pathways across the network, as discussed above. The comparison results suggest that bending-dependent disassembly acts as a form of mechanical quality control that limits the excessive accumulation of strongly deformed AFs, whereas repositioning to low-density regions acts as a form of spatial repair that replenishes gaps in the network. Our results suggest that, in living cells, these processes may act in coordination to limit excessive structural heterogeneity and help maintain cell-scale cortical tension as AFs are disassembled and repositioned.

When comparable amounts of myosin accumulated within the equatorial band, direct myosin localization and AF accumulation generated similar spatially averaged equatorward AF velocities, although some differences were observed in their latitudinal profiles. Here, the amount of bound myosin generating contractile force can be regarded as an indicator of contractile activity. Within the conditions examined here, this result suggests that comparable cell-scale cortical flows can arise at similar levels of equatorial contractile activity, regardless of whether myosin localizes directly to the band or through AF accumulation. Previous active-gel studies have shown that spatial gradients in actomyosin contractility drive cortical flow (Salbreux et al., 2009; Mayer et al., 2010). In these descriptions, cortical viscosity transmits flow away from the contractile region, whereas external friction attenuates it, so that the distance over which flow propagates is determined by the ratio of viscosity to friction. A filament-scale model has also shown that filament turnover and cross-link friction regulate stress dissipation and effective viscosity, thereby influencing cortical flow (McFadden et al., 2017). Although in-plane cortical viscosity is not explicitly represented in our model, contractile forces generated by myosin are transmitted as elastic stresses through the connected AF network, and these stresses may be relaxed through AF turnover and the binding and unbinding of network components. At timescales longer than the stress-relaxation timescale, this stress transmission and relaxation may behave as an effective viscous response and contribute to transmitting flow from the equatorial band, where contractile activity is high, to more distant regions.

Under AF accumulation, continuous high-tension structures formed along the band and high tension was concentrated within the equatorial band, whereas high-tension paths outside the band were relatively few and sparse. In contrast, under direct myosin localization, the branching of the high-tension network increased within the equatorial band, and high-tension paths were also broadly distributed outside the band up to mid-latitudes (Fig. 8(A)). In active-gel descriptions, cortical tension is expressed as the sum of active contractile stress and viscous stress, and its spatial gradient is balanced by external friction to generate flow (Mayer et al., 2010). Within this framework, tension within the equatorial band mainly reflects active contraction, whereas tension outside the band reflects not only basal contractile stress but also stress associated with meridional stretching of the cortex as it is drawn toward the equator. Consistent with this view, the meridional tension *τ*_*θθ*_ outside the band increased toward the equator under both conditions (Fig. 8(B)). In our model, the forces acting on each node are balanced by external friction; therefore, upon coarse-graining, the gradient in *τ*_*θθ*_ can be associated with the friction accompanying equatorward flow. Under direct myosin localization, *τ*_*θθ*_ decreased gradually over a broad range of latitudes, corresponding to the high-tension paths that extended to mid-latitudes. This suggests that a broader region of the cortex was stretched while being drawn toward the equator, consistent with the flow profile in which velocity was maintained up to higher latitudes (Fig. 7(B)). In contrast, under AF accumulation, the decrease in *τ*_*θθ*_ was concentrated near the equatorial band, and *τ*_*θθ*_ was nearly flat at higher latitudes. This is likely because AFs and force-generating myosin were concentrated within the equatorial band and AFs outside the band were depleted, thereby confining both the meridional-tension gradient and the flow to the vicinity of the equator. These results suggest that, even at similar levels of equatorial contractile activity, the mechanism of AF localization alters the structure of the network that bears contractile forces, and that this difference is reflected in the spatial distribution of flow through the latitudinal distribution of tension.

## 5. Limitations and future directions

The present model has several limitations. First, AF turnover was coarse-grained as the stochastic disassembly and subsequent relocation of entire filaments while maintaining a constant total number of AFs. In cells, the formation, remodeling, and turnover of cortical AFs involve multiple molecular processes, including nucleation, polymerization, depolymerization, severing, and branching, which alter filament number, length, and network architecture (Blanchoin et al., 2014). Because these processes were not explicitly represented, the present results identify the mechanical roles of effective AF removal and replenishment rules but do not directly describe the molecular kinetics or material fluxes underlying cortical turnover. Moreover, the mechanical dependence of AF severing varies with the mode of loading, and tensile loading has been reported to suppress cofilin binding and severing (Hayakawa et al., 2011). Thus, the bending-dependent disassembly rule in the present model should be regarded as an effective rule for removing buckled or strongly bent AFs and does not assume that all modes of mechanical loading promote AF disassembly. Explicit incorporation of these molecular processes and their mechanical dependence would enable the relationship between molecular-scale turnover dynamics and the maintenance of cell-scale cortical tension to be examined.

Second, the plasma membrane was represented as a prescribed spherical boundary, and cortical motion normal to the surface was constrained. Reconstituted and theoretical studies have shown that membrane tension can induce F-actin reorganization and flow, whereas cortical active tension and membrane–cortex friction can regulate cortical flow and bleb formation (Sakamoto et al., 2023; Sakamoto and Murrell, 2024). Consequently, the present model cannot capture the reciprocal coupling through which cortical tension and flow deform the membrane and changes in membrane shape and mechanical state feed back on cortical organization and force transmission. Coupling the present actomyosin cortex model to a deformable membrane model will be required to investigate these reciprocal dynamics.

Finally, the density-dependent AF relocation and the two myosin-localization mechanisms were introduced phenomenologically without explicitly modeling the associated biochemical regulators. In cells, mutual feedback between Rho and F-actin and reaction circuits involving Ect2 and RGA-3/4 contribute to the formation and maintenance of cortical patterns (Bement et al., 2015; Michaud et al., 2022). Furthermore, equatorial accumulation of AFs and myosin can involve at least partly distinct pathways, including cortical-flow-dependent transport, local assembly, and direct recruitment from the cytoplasm (Zhou and Wang, 2008; He et al., 2016). These cellular regulatory pathways do not correspond one-to-one to the localization mechanisms prescribed in the present model. Accordingly, the present comparisons identify the mechanical consequences of the prescribed spatial regulation but cannot establish how these patterns emerge or are dynamically maintained in cells. Future models that couple the discrete cortical network to reaction–diffusion dynamics of regulatory molecules would allow the feedback among biochemical signaling, network remodeling, tension generation, and cortical flow to be examined.

## 6. Conclusion

We investigated how filament-scale mechanics, turnover, and spatial reorganization collectively organize and maintain cortical tension and flow at the cellular scale, using a cellular-scale discrete model that explicitly represents AFs, CLs, and myosin minifilaments. The main conclusions are as follows:

- Cellular-scale cortical tension is organized by tension-bearing structures whose formation depends not only on component abundance but also on filament-scale mechanical loading, AF orientation, and network connectivity.
- The characteristic time scale of local tension-pattern reorganization was closest to the persistence time of myosin, suggesting that myosin dynamics may contribute to rapid force redistribution within the cortical network.
- AF relocation to low-density regions and bending-dependent disassembly play complementary roles in maintaining the cortical AF network by replenishing spatial gaps and limiting the accumulation of strongly deformed AFs.
- Direct myosin localization and AF accumulation generate comparable overall equatorward cortical flows, while differences in tension-bearing network organization are reflected in latitudinal tension and flow profiles.

## Acknowledgments

This study was partly supported by JSPS KAKENHI (Grant Numbers JP24KJ1869 and JP25K22796). Numerical simulations were performed on the TSUBAME4.0 supercomputer at the Institute of Science Tokyo.

## References

Amiri, S., Muresan, C., Shang, X., Huet-Calderwood, C., Schwartz, M. A., Calderwood, D. A. and Murrell, M., Intra-cellular tension sensor reveals mechanical anisotropy of the actin cytoskeleton, Nature Communications, Vol. 14, No.1 (2023), p.8011. Number: 1, DOI:10.1038/s41467-023-43612-5.

Bement, W. M., Leda, M., Moe, A. M., Kita, A. M., Larson, M. E., Golding, A. E., Pfeuti, C., Su, K.-C., Miller, A. L., Goryachev, A. B. and von Dassow, G., Activator–inhibitor coupling between Rho signalling and actin assembly makes the cell cortex an excitable medium, Nature Cell Biology, Vol. 17 (2015), pp.1471–1483, DOI:10.1038/ncb3251.

Blanchoin, L., Boujemaa-Paterski, R., Sykes, C. and Plastino, J., Actin Dynamics, Architecture, and Mechanics in Cell Motility, Physiological reviews, Vol. 94 (2014), pp.235–63, DOI:10.1152/physrev.00018.2013.

Chinthalapudi, K. and Heissler, S. M., Structure, regulation, and mechanisms of nonmuscle myosin-2, Cellular and Molecular Life Sciences, Vol. 81, No.1 (2024), p.263, DOI:10.1007/s00018-024-05264-6.

Chugh, P., Clark, A. G., Smith, M. B., Cassani, D. A. D., Dierkes, K., Ragab, A., Roux, P. P., Charras, G., Salbreux, G. and Paluch, E. K., Actin cortex architecture regulates cell surface tension, Nature Cell Biology, Vol. 19, No.6 (2017), pp.689–697. Number: 6, DOI:10.1038/ncb3525.

Clausen, M. P., Colin-York, H., Schneider, F., Eggeling, C. and Fritzsche, M., Dissecting the actin cortex density and membrane-cortex distance in living cells by super-resolution microscopy, Journal of Physics D: Applied Physics, Vol. 50, No.6 (2017), p.064002, DOI:10.1088/1361-6463/aa52a1.

Hayakawa, K., Tatsumi, H. and Sokabe, M., Actin filaments function as a tension sensor by tension-dependent binding of cofilin to the filament, Journal of Cell Biology, Vol. 195, No.5 (2011), pp.721–727, DOI:10.1083/jcb.201102039.

He, B., Martin, A. C. and Wieschaus, E., Flow-dependent myosin recruitment during Drosophila cellularization requires zygotic dunk activity, Development, Vol. 143, No.13 (2016), pp.2417–2430, DOI:10.1242/dev.131334.

Hernández-del Valle, M., Valencia-Expósito, A., López-Izquierdo, A., Casanova-Ferrer, P., Tarazona, P., Martín-Bermudo, M. D. and Míguez, D. G., A coarse-grained approach to model the dynamics of the actomyosin cortex, BMC Biology, Vol. 20, No.1 (2022), p.90, DOI:10.1186/s12915-022-01279-2.

Kelkar, M., Bohec, P. and Charras, G., Mechanics of the cellular actin cortex: From signalling to shape change, Current Opinion in Cell Biology, Vol. 66 (2020), pp.69–78, DOI:10.1016/j.ceb.2020.05.008.

Kim, J. H. and Kim, T., The mechanism of dynamic steady states in lamellipodia, PLOS Computational Biology, Vol. 21, No.10 (2025), p.e1013572, DOI:10.1371/journal.pcbi.1013572.

Kim, T., Hwang, W., Lee, H. and Kamm, R. D., Computational Analysis of Viscoelastic Properties of Crosslinked Actin Networks, PLOS Computational Biology, Vol. 5, No.7 (2009), p.e1000439, DOI:10.1371/journal.pcbi.1000439.

Kojima, H., Ishijima, A. and Yanagida, T., Direct measurement of stiffness of single actin filaments with and without tropomyosin by in vitro nanomanipulation., Proceedings of the National Academy of Sciences, Vol. 91, No.26 (1994), pp.12962–12966, DOI:10.1073/pnas.91.26.12962.

Krishna, A., Savinov, M., Ierushalmi, N., Mogilner, A. and Keren, K., Size-dependent transition from steady contraction to waves in actomyosin networks with turnover, Nature Physics, Vol. 20, No.1 (2024), pp.123–134, DOI:10.1038/s41567-023-02271-5.

Lenz, M., Geometrical Origins of Contractility in Disordered Actomyosin Networks, Physical Review X, Vol. 4, No.4 (2014), DOI:10.1103/PhysRevX.4.041002.

Maxian, O., Peláez, R. P., Mogilner, A. and Donev, A., Simulations of dynamically cross-linked actin networks: Morphology, rheology, and hydrodynamic interactions, PLOS Computational Biology, Vol. 17, No.12 (2021), p.e1009240, DOI:10.1371/journal.pcbi.1009240.

Maxian, O., Dinner, A. R. and Munro, E., Actin network heterogeneity tunes activator ‒ inhibitor dynamics at the cell cortex, Proceedings of the National Academy of Sciences, Vol. 122, No.50 (2025), p.e2520485122, DOI:10.1073/pnas.2520485122.

Mayer, M., Depken, M., Bois, J. S., Jülicher, F. and Grill, S. W., Anisotropies in cortical tension reveal the physical basis of polarizing cortical flows, Nature, Vol. 467, No.7315 (2010), pp.617–621, DOI:10.1038/nature09376.

McCall, P. M., MacKintosh, F. C., Kovar, D. R. and Gardel, M. L., Cofilin drives rapid turnover and fluidization of entangled F-actin, Proceedings of the National Academy of Sciences, Vol. 116, No.26 (2019), pp.12629–12637, DOI:10.1073/pnas.1818808116.

McFadden, W. M., McCall, P. M., Gardel, M. L. and Munro, E. M., Filament turnover tunes both force generation and dissipation to control long-range flows in a model actomyosin cortex, PLOS Computational Biology, Vol. 13, No.12 (2017), p.e1005811, DOI:10.1371/journal.pcbi.1005811.

Michaud, A., Leda, M., Swider, Z. T., Kim, S., He, J., Landino, J., Valley, J. R., Huisken, J., Goryachev, A. B., von Dassow, G. and Bement, W. M., A versatile cortical pattern-forming circuit based on Rho, F-actin, Ect2, and RGA-3/4, Journal of Cell Biology, Vol. 221, No.8 (2022), p.e202203017, DOI:10.1083/jcb.202203017.

Murrell, M., Oakes, P. W., Lenz, M. and Gardel, M. L., Forcing cells into shape: the mechanics of actomyosin contractility, Nature Reviews Molecular Cell Biology, Vol. 16, No.8 (2015), pp.486–498, DOI:10.1038/nrm4012.

Murrell, M. P. and Gardel, M. L., F-actin buckling coordinates contractility and severing in a biomimetic actomyosin cortex, Proceedings of the National Academy of Sciences, Vol. 109, No.51 (2012), pp.20820–20825, DOI:10.1073/pnas.1214753109.

Popov, K., Komianos, J. and Papoian, G. A., MEDYAN: Mechanochemical Simulations of Contraction and Polarity Alignment in Actomyosin Networks, PLOS Computational Biology, Vol. 12, No.4 (2016), p.e1004877, DOI:10.1371/journal.pcbi.1004877.

Ramanathan, S. P., Helenius, J., Stewart, M. P., Cattin, C. J., Hyman, A. A. and Muller, D. J., Cdk1-dependent mitotic enrichment of cortical myosin II promotes cell rounding against confinement, Nature Cell Biology, Vol. 17, No.2 (2015), pp.148–159. Number: 2, DOI:10.1038/ncb3098.

Reymann, A.-C., Staniscia, F., Erzberger, A., Salbreux, G. and Grill, S. W., Cortical flow aligns actin filaments to form a furrow, eLife, Vol. 5 (2016), p.e17807, DOI:10.7554/eLife.17807.

Rutkowski, D. M. and Vavylonis, D., Discrete mechanical model of lamellipodial actin network implements molecular clutch mechanism and generates arcs and microspikes, PLoS Computational Biology, Vol. 17, No.10 (2021), p.e1009506, DOI:10.1371/journal.pcbi.1009506.

Sakamoto, R., Banerjee, D. S., Yadav, V., Chen, S., Gardel, M., Sykes, C., Banerjee, S. and Murrell, M. P., Membrane tension induces F-actin reorganization and flow in a biomimetic model cortex, Communications Biology, Vol. 6, No.1 (2023), pp.1–10. Number: 1, DOI:10.1038/s42003-023-04684-7.

Sakamoto, R. and Murrell, M. P., Active tension and membrane friction mediate cortical flows and blebbing in a model actomyosin cortex, Physical Review Research, Vol. 6, No.3 (2024), p.033024, DOI:10.1103/PhysRevResearch.6.033024.

Salbreux, G., Charras, G. and Paluch, E., Actin cortex mechanics and cellular morphogenesis, Trends in Cell Biology, Vol. 22, No.10 (2012), pp.536–545, DOI:10.1016/j.tcb.2012.07.001.

Salbreux, G., Prost, J. and Joanny, J. F., Hydrodynamics of Cellular Cortical Flows and the Formation of Contractile Rings, Physical Review Letters, Vol. 103, No.5 (2009), p.058102, DOI:10.1103/PhysRevLett.103.058102.

Schramm, A. C., Hocky, G. M., Voth, G. A., Blanchoin, L., Martiel, J.-L. and De La Cruz, E. M., Actin Filament Strain Promotes Severing and Cofilin Dissociation, Biophysical Journal, Vol. 112, No.12 (2017), pp.2624–2633, DOI:10.1016/j.bpj.2017.05.016.

Schramm, A. C., Hocky, G. M., Voth, G. A., Martiel, J.-L. and De La Cruz, E. M., Plastic Deformation and Fragmentation of Strained Actin Filaments, Biophysical Journal, Vol. 117, No.3 (2019), pp.453–463, DOI:10.1016/j.bpj.2019.06.018.

Spira, F., Cuylen-Haering, S., Mehta, S., Samwer, M., Reversat, A., Verma, A., Oldenbourg, R., Sixt, M. and Gerlich, D. W., Cytokinesis in vertebrate cells initiates by contraction of an equatorial actomyosin network composed of randomly oriented filaments, eLife, Vol. 6 (2017), p.e30867, DOI:10.7554/eLife.30867.

Staddon, M. F., Munro, E. M. and Banerjee, S., Pulsatile contractions and pattern formation in excitable actomyosin cortex, PLOS Computational Biology, Vol. 18, No.3 (2022), p.e1009981, DOI:10.1371/journal.pcbi.1009981.

Svitkina, T. M., Actin Cell Cortex: Structure and Molecular Organization, Trends in Cell Biology, Vol. 30, No.7 (2020), pp.556–565, DOI:10.1016/j.tcb.2020.03.005.

Tam, A. K. Y., Mogilner, A. and Oelz, D. B., Nonuniform filament turnover, contractility, and bundle formation in disordered actomyosin networks, Biophysical Journal, Vol. 125, No.8 (2026), pp.1867–1879, DOI:10.1016/j.bpj.2026.02.032.

Truong Quang, B. A., Peters, R., Cassani, D. A. D., Chugh, P., Clark, A. G., Agnew, M., Charras, G. and Paluch, E. K., Extent of myosin penetration within the actin cortex regulates cell surface mechanics, Nature Communications, Vol. 12, No.1 (2021), p.6511. Number: 1, DOI:10.1038/s41467-021-26611-2.

Vogel, S. K., Petrasek, Z., Heinemann, F. and Schwille, P., Myosin motors fragment and compact membrane-bound actin filaments, eLife, Vol. 2 (2013), p.e00116, DOI:10.7554/eLife.00116.

Yu, Q., Li, J., Murrell, M. P. and Kim, T., Balance between Force Generation and Relaxation Leads to Pulsed Contraction of Actomyosin Networks, Biophysical Journal, Vol. 115, No.10 (2018), pp.2003–2013, DOI:10.1016/j.bpj.2018.10.008.

Zhou, M. and Wang, Y.-L., Distinct Pathways for the Early Recruitment of Myosin II and Actin to the Cytokinetic Furrow, Molecular Biology of the Cell, Vol. 19, No.1 (2008), pp.318–326, DOI:10.1091/mbc.E07-08-0783.

